# *PyCycleBio:* modelling non-sinusoidal-oscillator systems in temporal biology

**DOI:** 10.1101/2025.04.30.651403

**Authors:** Alexander R. Bennett, George Birchenough, Daniel Bojar

**Affiliations:** Department of Medical Biochemistry, Institute of Biomedicine, University of Gothenburg, 41390 Gothenburg, Sweden; Department of Chemistry and Molecular Biology, Institute of Biomedicine, University of Gothenburg, 41390 Gothenburg, Sweden; Wallenberg Centre for Molecular and Translational Medicine, University of Gothenburg, 41390 Gothenburg, Sweden

## Abstract

Protein, mRNA, and metabolite abundances can exhibit rhythmic dynamics, such as during the day/night cycle. Leading bioinformatics platforms for identifying biological rhythms often utilise single-component models of the harmonic oscillator equation, or multi-component models based upon the Cosinor framework. These approaches offer distinct advantages: modelling either temporally-resolved regulatory behaviour via the extended harmonic oscillator equation, or complex rhythmic patterns in the case of Cosinor. Here, we have developed a new platform to combine the advantages of these two approaches. *PyCycleBio* utilises bounded-multi-component models and modulus operators alongside the harmonic oscillator equation, to model a diverse and interpretable array of rhythmic behaviours, including the regulation of temporal dynamics via amplitude coefficients. We demonstrate increased sensitivity and functionality of *PyCycleBio* compared to other analytical frameworks, and uncover new relationships between data modalities or sampling conditions with the qualities of rhythmic behaviours from biological datasets— including transcriptomics, proteomics, and metabolomics. We envision that this new approach for disentangling complicated temporal regulation of biomolecules will advance chronobiology and our understanding of physiology. *PyCycleBio* is available at: https://github.com/Glycocalex/PyCycleBio, and the Python package is available to install at: https://pypi.org/project/pycyclebio/. *PyCycleBio* can also be used at https://colab.research.google.com/github/Glycocalex/PyCycleBio/blob/main/PyCycleBio.ipynb with no installations necessary.

## 1 Introduction

Many behavioural and physiological processes exhibit rhythmic dynamics, which are adaptations to regular phenomena such as the day-night cycle. In recent decades, the molecular basis for many of these processes has been elucidated (Partch, *et al*. 2014). Circadian rhythms and the cell cycle are among the better-characterised biological rhythms. Many chronobiological phenomena can be accurately described with a sinusoidal curve—the simplest model providing stable oscillatory behaviour—and sinusoidal oscillations are commonly used to identify rhythmic biological molecules. However it has long been noted that many rhythmic processes violate sinusoidal assumption (Flesia, *et al*. 2022). Theoretical work has demonstrated that non-sinusoidal oscillations are for instance required for temperature compensation, an important and widely conserved phenomenon where the rate of circadian processes is maintained while temperature changes (Gibo, Kurosawa, 2019). Behavioural data is also well appreciated to exhibit non-sinusoidal rhythms in many cases (Cichewicz, Hirsh, 2018), and many other biological rhythms exhibit non-sinusoidal dynamics, such as sexual (Zheng, *et al*. 2024) or seasonal cycles (Nagano, *et al*. 2019). These biological rhythms remain relatively underexplored, compared to circadian or cell-cycle phenomena, and improved bioinformatics support for complex-rhythm analysis may facilitate future studies in these areas. Additionally, the often-observed low concordance between rhythmicity among corresponding transcripts and proteins suggests that multiple regulatory mechanisms influence observable dynamics (de Los Santos, *et al*. 2020, Bignon, *et al*. 2023). Since current methods usually cannot easily model these phenomena, we currently have an incomplete understanding of how biological rhythms manifest, which is partly due to a lack of non-sinusoidal modelling techniques.

Current models already provide meaningful insights into the dynamics of regulatory elements that give rise to temporal patterns. Temporal regulation of biological processes is important in homeostasis and disease, and an improved understanding of temporal rhythms facilitates identification of key regulatory pathways underpinning these behaviours. Parametric models also make it possible to deduce the relative periodicities and amplitudes of linked oscillatory elements, informing further investigation.

Since its introduction to the field, the Jonckheere-Terpstra-Kendall (JTK) algorithm (Hughes, *et al*. 2010) has been widely adopted as an effective and approachable framework for describing sinusoidal expression patterns in transcriptomics or proteomics data, using a non-parametric mapping of the harmonic oscillator equation. More recently, de Los Santos, *et al*. (2020) have expanded upon the framework of ‘JTK’ by introducing the *Extended Circadian Harmonic Oscillator* (ECHO), a statistical framework to parametrically model patterns of expression exhibiting linear increases or decreases in amplitude over the temporal period of analysis. This was an important development that modelled the regulation of oscillatory dynamics and highlighted, for example, the differential regulation of rhythmicity in corresponding elements of the transcriptome and proteome of *Neurospora crassa*. Additionally, the ‘Cosinor’ model has long been used for biological rhythm analysis (Halberg, *et al*. 1967), and was recently developed into a Python package (Moškon, 2020). Cosinor utilises a multi-oscillator system for parameterising temporal data and, critically, modern Cosinor analysis can model non-sinusoidal rhythms. Thus, both Cosinor and ECHO facilitate the understanding of more complex regulatory mechanisms, and together they represent the current gold-standard for statistical analysis of biorhythmic data.

Presently however, the advantages offered by both of these platforms remain distinct. The multi-component nature of Cosinor analysis makes it difficult to implement alongside a single-component harmonic-oscillator model, and although Cosinor can parametrise complex waveforms, identifying regulatory elements that give rise to observed oscillations is challenging. Cosinor frameworks are also not equipped to employ variable numbers of oscillatory components within an analysis. Further, including many components—while producing accurate parameterisations—can result in biological signals that are difficult to interpret and risk being distorted by noise.

For these reasons, we have developed *PyCycleBio*, an analytical framework that couples the modulatory coefficient of the extended harmonic oscillator equation with bounded-multi-component waveforms, inspired by Cosinor. Through *PyCycleBio*, we combine the best elements of both approaches into a functional and easy-to-use toolset which models sinusoidal and complex oscillations in a data-driven manner. This results in parameterisations that are more functional and interpretable than intensive algorithms, producing bespoke fits on a per-molecule basis. Using a least-squares method for fitting model predictions to observed data also makes *PyCycleBio* robust and applicable to datasets with a non-uniform sampling regime. Here, we outline the theory behind our model and use it to identify non-sinusoidal rhythms from published transcriptomics, proteomics, and metabolomics datasets, which are missed by current analysis platforms. Overall, we present *PyCycleBio* as a new foundation for data-driven chronobiology.

## 2 Methodology

### 2.1 Model definition

*PyCycleBio* is capable of modelling: (i) dampened or forced sine waves, using the extended harmonic oscillator equation; (ii) ‘pseudo-square waves’, approximating the digital waveform with two sinusoidal components; (iii) ‘cycloid waves’, where sinusoidal expression is modulated by a second sine wave of double amplitude but half-periodicity; and (iv) ‘transients’, where a modulo operator is employed to produce periodic increases in magnitude from a stable baseline.

The extended harmonic oscillator equation is employed in the state-of-the-art bioinformatics platform ECHO (de Los Santos, *et al*. 2020), and we have utilised the same equation for the parameterisation of sinusoidal oscillations in *PyCycleBio* :

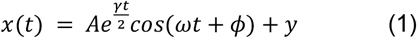

where x(t) is the amplitude at time *t*, A is the initial amplitude (*t*=0), γ is the amplitude change coefficient, ω is the frequency of oscillation, ϕ is the phase shift, and y is the equilibrium value. We have expanded upon (1) with a pair of equations that each feature two sinusoidal terms: The ‘pseudo-square wave’ equation approximates the digital square waveform using two oscillators as follows (Smith, S. 1997):

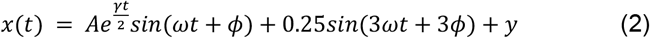

In equation (2), the carrier waveform is modulated using the sum of a second sinusoidal oscillator with one-third the periodicity and half the amplitude of the carrier. The ‘pseudo-cycloid wave’ is another revision of (1), and employs two cosine components to approximate the dynamics of a cycloid wave:

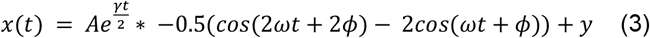

Finally we have utilised a modulator to apply a Gaussian function at specific intervals, producing transient expression impulses from a stable baseline:

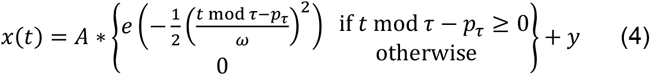

### 2.2 Model selection

The initial stage of modelling biological molecule expression is to determine the optimal parameterisation of each equation, producing the closest-fit to the measured temporal data. Our algorithm employs the *optimize.curve_fit* function from SciPy (version 1.11.1; Virtanen, *et al*. 2020) to produce an optimal parameterisation via non-linear least square fitting to iteratively manipulate starting parameters within bounds. We note that SciPy utilises Fortran for this, which offers substantial performance benefits compared to pure Python. As a result, *PyCycleBio* analyses are substantially faster than comparable platforms, with most analyses around 70-80x quicker than parametric alternatives **(S.fig. 1A)**.

These parameters are then used to produce fitted values, and the residuals between fitted and experimentally-measured values are computed. Residuals are squared, so deviations above and below fitted values contribute equally to the residual magnitude. Once optimal parameters and residuals have been calculated for all models, the model with the lowest sum of residuals is determined to be optimal.

The final stage of modelling is to determine how robustly the optimal model describes the experimentally-measured values. For this, Kendall’s tau algorithm is employed, from the implementation available in the *stats* module of SciPy (version 1.11.1; Virtanen, *et al*. 2020), statistically determining how accurately fitted values represent experimental values. Model outputs are provided alongside the p-value of the optimal model. We additionally report the Benjamini-Hochberg corrected p-values.

### 2.3 Datasets

Datasets analysed in this work have been sourced from the academic literature, by processing supplementary tables. Synthetic data was generated from the *Timetrial* R package (v1.0), at variance levels of 0.1 and 0.4 (Ness-Cohn, *et al*. 2020). In general, collected datasets had been already normalised and so no alterations to the original data were performed, unless otherwise specified. For comparisons between *PyCycleBio* and other methods, the software and version numbers were as follows: JTK_CYCLE v3.1 (R), ECHO v4.0 (R), CosinorPy v3.1 (Python). R version 4.3.0 and Python 3.11 were used for all analyses in this work. In all, we used supplementary tables from: Acosta-Rodrigez, et al. 2022; Aviram, et al. 2021; Benegiamo, et al. 2018; Bignon, et al. 2023 and Cichewicz and Hirsh, 2018.

## 3 Results

### 3.1 PyCycleBio models distinct rhythmic waveforms

*PyCycleBio* is able to parameterise: sinusoidal, cycloid, square, and transient patterns of expression, with amplitude coefficients to indicate active regulatory processes. Idealised versions of these waveforms, as well as example transcriptomics data exhibiting these forms of rhythmic expression are visualised in **Fig. 1A**, with examples of regulatory classifications in **Fig. 1B** (Bignon, *et al*. 2023). We note that behavioural data in particular is poorly modelled with sinusoidal waves. Locomotor activity data from *Drosophila melanogaster*, recorded by the *Drosophila* Activity Monitor system (Cichewicz and Hirsh, 2018), was analysed with *PyCycleBio* and confirmed a square waveform to be more suited for parameterisation than a sine wave, corroborated by a lower RMSE **(Fig. 1C)**.

**Figure 1:**
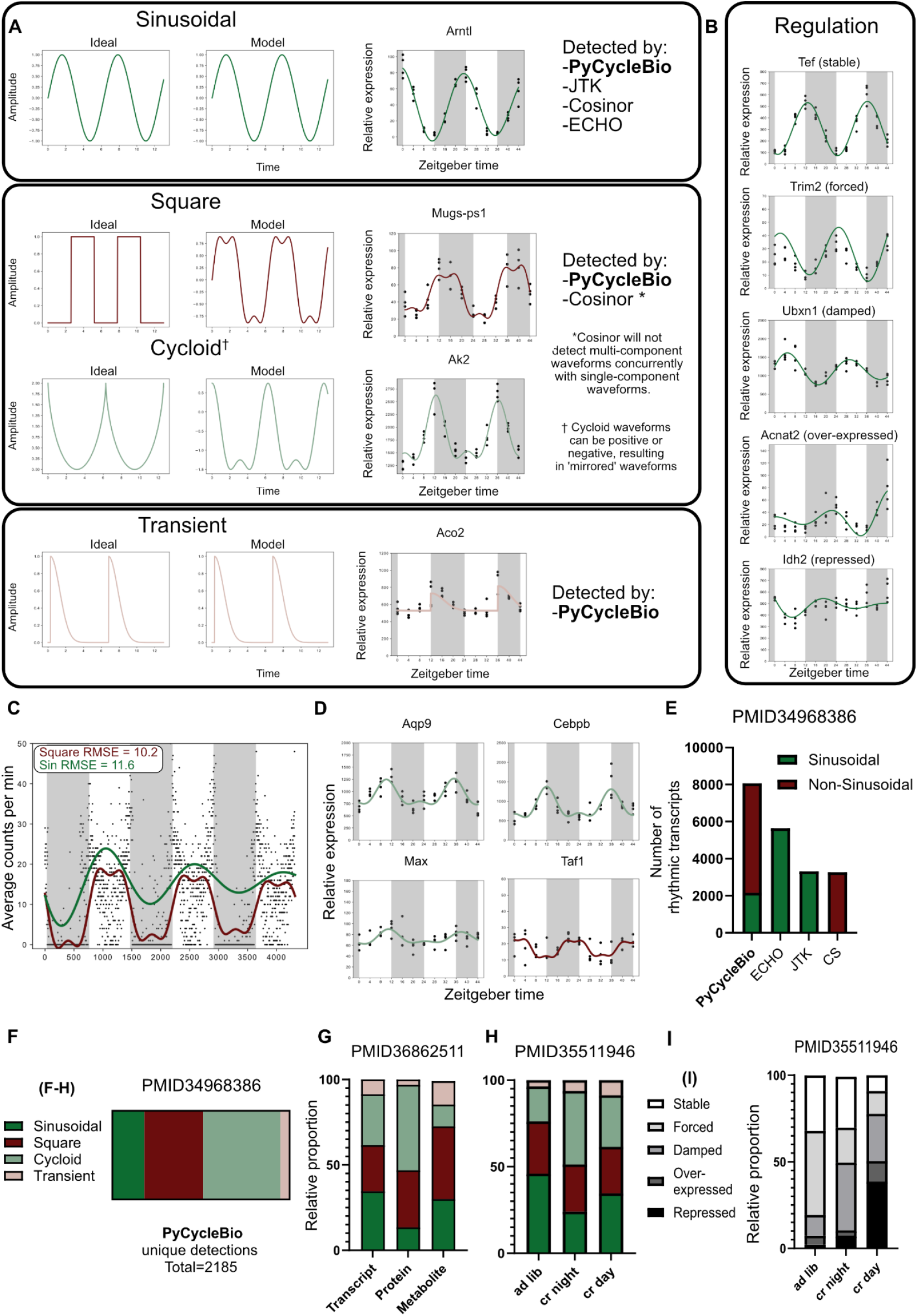
Waveform characterisation, model performance, and interpretation. **(A)** Plots visualising an ideal waveform, our modelled representation, and an example transcript corresponding to each type of waveform (Aviram, *et al*. 2021). **(B)** Examples of regulatory classifications based upon *γ* in equations (1), (2), and (3) (Aviram, *et al*. 2021). Transcriptomics data are presented as mean +/-SD of z-score normalised expression data. **(C)** An example of the optimal square and sinusoidal models produced to fit behavioural data (data adapted from Cichewicz and Hirsh, 2018). **(D)** Expression of *Aqp9* and transcription factors *Cebpb, Max*, and *Taf1* (Aviram, *et al*. 2021). **(E)** The total number of rhythmic transcripts in a transcriptomics dataset (Bignon, *et al*. 2023), as identified by *PyCycleBio* and other common analytical platforms. **(F)** Distribution of assigned models in transcripts (Bignon, *et al*. 2023) identified as rhythmic by *PyCycleBio* and not by *ECHO*. **(G)** Distribution of models in three tissue-matched ‘omics modalities, generated from murine renal tissue (Bignon, *et al*. 2023). **(H)** Distributions of models and **(H)** regulatory behaviour of oscillating molecules, within the hepatic transcriptome of mice fed *ad-libitum*, for two hours during the night or for two hours during the day (Acosta-Rodrigez, *et al*. 2022).

Transcriptomics data from murine liver tissue demonstrates how *PyCycleBio* can facilitate interpretation of the regulatory insights giving rise to observed regulatory dynamics (Aviram, *et al*. 2021). For example, the cycloid dynamics observed in *Aqp9* expression **(Fig. 1D)** are modelled by equation (3), which describes the additive effects of two oscillatory elements: one of which is circadian and the other ultradian, with a period of approximately 12 hours. This is corroborated by the observed expression of transcription factors *Cebpb, Max*, and *Taf1. Aqp9* has two promoter regions: *Cebpb* binds one promoter and has a 24-hour rhythm at the transcript level; the second *Aqp9* promoter is bound by both *Max* and *Taf1*, which oscillate with anti-phase 24-hour rhythms, suggesting that one of these factors is active on the promoter every 12 hours. Together, the dynamics of these three transcription factors activating two promoter regions could explain the observed expression dynamics of *Aqp9*.

### 3.2 PyCycleBio outperforms other temporal bioinformatics platforms

Thanks to our highly optimized implementation, *PyCycleBio* also offers substantially improved performance, typically on the order of 70-80x faster, compared to other analytical platforms when analysing both synthetic and biological datasets **(S.fig. 1A)**. Utilising synthetic data from Ness-Cohn, *et al*. (2020), we compared the accuracy of *PyCycleBio* to ECHO and Cosinor using receiver-operating-characteristic curves. Curves were generated for synthetic data with noise (variance) levels of 0.1x and 0.4x total signal amplitude **(S.fig. 1B)**. Overall, we observed that *PyCycleBio* performed best out of the three platforms tested at both noise levels, as indicated by ROC area under the curve values of 0.99 and 0.97 at noise levels of 0.1 and 0.4, respectively. We further note that the synthetic dataset contained only sinusoidal data models, and did not even leverage *PyCycleBio’s* primary advantage of detecting non-sinusoidal signals.

### 3.3 The majority of unique rhythms detected by PyCycleBio are non-sinusoidal

We find *PyCycleBio* to be more sensitive in detecting oscillatory behaviours than other commonly-used bioinformatics packages **(Fig. 1E)**: 5,526 rhythmic molecules were detected by both *PyCycleBio* and ECHO, 2,544 by *PyCycleBio* alone, and only 121 by ECHO alone **(S.Fig. 1C)**. Although increased detection is an attractive selling-point for analytical algorithms, minimising false-positives is essential too. *PyCycleBio* thus uses a false-discovery-control procedure using the Benjamini-Hochberg correction. Additionally, we note that, although the total number of rhythmic molecules was generally increased in a *PyCycleBio* analysis compared to other frameworks, the number of sinusoidal oscillations was reduced, suggesting many rhythmic molecules were better classified using our non-sinusoidal models. Reanalysing the dataset generated by Menalla, *et al*. (2021) revealed that only 16.5% of the oscillatory molecules detected only by *PyCycleBio*, but not ECHO, were best modelled using the harmonic oscillator equation **(Fig. 1F)**. This indicated that, although *PyCycleBio* detected more rhythmic transcripts than ECHO, this was mainly due to the identification of additional non-sinusoidal behaviours, which ECHO was not designed to detect.

### 3.4 Data modalities and sampling conditions influence rhythm type distribution

As we have shown above, surprisingly many biomolecules exhibit non-sinusoidal dynamics. We thus hypothesised that different data modalities or sampling conditions may exhibit different distribution of biorhythmic models. To investigate this, we re-analysed previously published datasets from circadian studies with high sampling frequencies. As one example, Bignon, *et al*. (2023) compared the transcriptome, proteome, and metabolome of murine renal tissue **(Fig. 1G)**. Analysis of each dataset by *PyCycleBio* revealed different distributions of rhythm types between modalities. The transcriptome exhibited the greatest proportion of sinusoidal oscillations (34.6%), in contrast to the proteome, where only 13.5% of oscillatory proteins were best modelled by a sinusoidal oscillation. Interestingly, the metabolome had the highest proportion of transient molecules at 13.8%, whereas the transcriptome and proteome exhibited 8.7% and 3.1%, respectively.

Further, we utilised transcriptomics data generated by Acosta-Rodrigez, *et al*. (2022), in which mice were either fed *ad-libitum*, or calorie-restricted to two hours during the light or dark phases **(Fig. 1H)**. We observed shifts in the rhythmic content of the liver transcriptome following calorie-restriction, with sinusoidal oscillations being reduced from 45.9% in *ad-lib* fed mice to 24.1% in night-fed and 34.6% in day-fed mice. Additionally, the damping coefficient obtained with our analyses provided insights into regulatory behaviours of the observed rhythms, as 32.0% of rhythms observed in *ad-lib*-fed mice exhibited stable behaviour (not actively being up- or down-regulated during the sampled timeframe). Conversely, this was reduced to 29.4% of rhythms being stably expressed in night-fed animals, while in day-fed animals only 9.1% of rhythms were stable over the experimental time-course **(Fig. 1I)**. Restricted feeding during the night induced rhythmic expression in 4,122 transcripts, while 5,638 transcripts gained rhythmicity in day-fed animals. Overall, 2,019 transcripts gained rhythmicity in both restricted-feeding groups, indicating that the majority of rhythmic transcripts are unique to either condition. Further, approximately half of rhythmic transcripts from the *ad-lib*-fed mice lost rhythmicity after restricted feeding. Mice fed during the night (the active phase), were more similar to *ad-lib* mice in this than mice fed during the day (inactive phase). This suggests stronger disruptions to diurnal behaviours can manifest as increased heterogeneity in the regulation and form of rhythmic transcriptome content, detectable via *PyCycleBio*, which should be considered in experimental design and physiological outcomes.

## 4) Discussion

*PyCycleBio* couples the amplitude-change coefficient employed in ECHO (de Los Santos, *et al*. 2019) with the multi-component models of Cosinor (Cornelissen, 2014). We have validated the capacity of our approach to identify complex biological rhythms, demonstrating it to be as sensitive to sinusoidal rhythms as the current state-of-the-art **(S.fig. 1B)** as well as vastly more performant **(S.fig. 1A)**. We note that the majority of rhythms detected by *PyCycleBio* are non-sinusoidal, evidencing the importance of considering complex dynamics **(Fig. 1F)**. We observe that disrupting biological rhythms, for instance by calorie-restricting mice to a short window during the night or day, resulted in reduced sinusoidal content of the rhythmic molecular compartment **(Fig. 1H)** and exhibited a plethora of actively regulated rhythmic molecules **(Fig. 1G)**. We suggest this could reflect the dynamic control of larger physiological programs with sinusoidal oscillations, which yield more complex dynamics when perturbed. Additionally, the distribution of rhythmic models differed between data modalities. Transcriptomcs data appeared to have the highest sinusoidal content, while proteomics and metabolomics exhibited more complex dynamics **(Fig. 1G)**. Post-translational regulatory processes governing protein and metabolite abundance could offer a possible explanation for this observation, as many distinct factors can contribute to the observed complex dynamics in these instances. We also note differences in the distribution of rhythmic behaviours between datasets **(Fig. 1 G,H)**, which may reflect distinct peripheral clock behaviours in different tissues, highlighting the interesting new lines of inquiry our approach facilitates.

We also plan to expand the functionality of *PyCycleBio* in the future to perform differential rhythmicity analysis, as is facilitated by some current methods such as DryR (Weger, *et al*. 2021) and Circacompare (Parsons, *et al*. 2020).

We envision *PyCycleBio* to be useful for the circadian and chronobiological fields, aiding our understanding of complex temporal phenomena and the underlying regulatory mechanisms. Additionally, we note that most temporal bioinformatics platforms have been developed and deployed using R. Developed in Python as open-source software, *PyCycleBio* can leverage, and be integrated alongside, computational platforms from other fields, such as data science and machine learning. We are thus confident *PyCycleBio* will enable chronobiologists to benefit from exciting developments in computational biology and artificial intelligence in the future.

---

**Supplementary figure 1:**
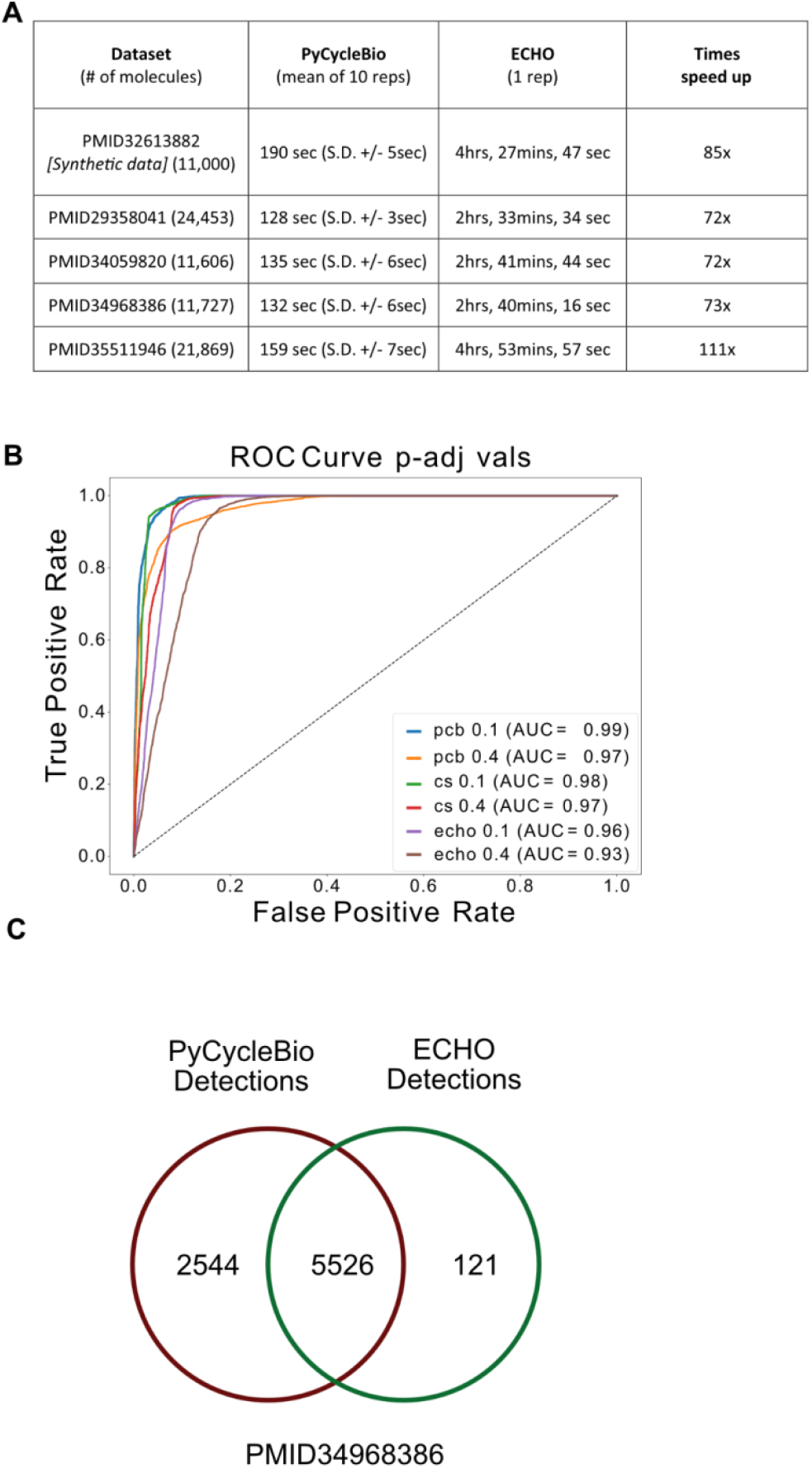
**(A)** Receiver-operating-characteristic (ROC) curves for the accuracy of three platforms: *PyCycleBio* (pcb), *Cosinor* (cs), and *ECHO* (echo), when analysing synthetic sinusoidal data with a noise ratio of 0.1x or 0.4x total amplitude (Ness-Cohn, 2020). **(B)** A comparison of the time required to analyse circadian datasets, between *PyCycleBio* and *ECHO* (de Los Santos, *et al*. 2020). **(C)** A Venn-diagram showing the number of common and unique rhythmic transcripts detected by *PyCycleBio* and ECHO (using data from Bignon, *et al*. 2023).

